# The activity of *engrailed* imaginal disc enhancers is modulated epigenetically by chromatin and autoregulation

**DOI:** 10.1101/2023.06.15.545191

**Authors:** Yuzhong Cheng, Fountane Chan, Judith A. Kassis

**Affiliations:** Division of Developmental Biology, Eunice Kennedy Shriver National Institute of Child Health and Human Development, National Institutes of Health, Bethesda, MD 20892

## Abstract

*engrailed* (*en*) encodes a homeodomain transcription factor crucial for the proper development of Drosophila embryos and adults. Like many developmental transcription factors, *en* expression is regulated by many enhancers, some of overlapping function, that drive expression in spatially and temporally restricted patterns. The *en* embryonic enhancers are located in discrete DNA fragments that can function correctly in small reporter transgenes. In contrast, the *en* imaginal disc enhancers (IDEs) do not function correctly in small reporter transgenes. En is expressed in the posterior compartment of wing imaginal disks; small IDE-reporter transgenes are expressed in the anterior compartment, the opposite of what is expected. Our data show that the En protein binds to *en* IDEs, and we suggest that En directly represses IDE function. We identified two *en* IDEs, ‘O’ and ‘S’. Deletion of either of these IDEs from a 79kb HA-en rescue transgene (*HAen79*) caused a loss-of-function *en* phenotype when the *HAen79* transgene was the sole source of En. In contrast, flies with a deletion of the same IDEs from the endogenous *en* gene had no phenotype, suggesting a resiliency not seen in the *HAen79* rescue transgene. Inserting a gypsy insulator in *HAen79* between *en* regulatory DNA and flanking sequences strengthened the activity of *HAen79*, giving better function in both the ON and OFF transcriptional states. Altogether our data show that the *en* IDEs stimulate expression in the entire imaginal disc, and that the ON/OFF state is set by epigenetic regulators. Further, the endogenous locus imparts a stability to *en* function not seen even in a large transgene, reflecting the importance of both positive and negative epigenetic influences that act over relatively large distances in chromatin.

**Author Summary:** Genes that control development are often used at different times and places in a developing embryo. Transcription of these important genes must be tightly regulated; therefore, these genes often have large arrays of regulatory DNA. In Drosophila, discrete fragments of DNA (enhancers) can be identified that turn genes on in patterns in the early embryo. In cells where the genes are transcriptionally ON, there are active modifications on chromatin, setting later enhancers in a transcription-permission environment. In cells where the genes are OFF, repressive chromatin marks keep later enhancers inactive. In this paper we studied two late enhancers of the Drosophila *en* gene. We show that the correct activity of these enhancers is dependent on being next to other, earlier acting *en* enhancers. Our data also show that En can repress its own expression, likely directly by acting on these late enhancers. The chromatin-regulated activity of these *en* late enhancers is similar to what was described for a late enhancer of another Drosophila developmental gene, *Ubx*. We suggest that this mode of regulation is likely to be important for many late-acting developmental enhancers in many different organisms.

## Introduction

Developmentally important transcription factors are expressed in spatially and temporally restricted patterns in the precursors of many different cell types. These complex gene expression patterns are generated by a large number of enhancers, traditionally defined by their abilities to stimulate patterned gene expression in transgenes (reviewed in Small and Arnosti 2020). Many developmental genes have so-called “shadow enhancers”; that is, more than one enhancer that can drive transcription in a similar pattern. Enhancers with overlapping functions are thought to impart robustness to transcription of these important genes (Frankel et al. 2010; Perry et al. 2010; Osterwalder et al 2018; reviewed in Kvon et al. 2021). In addition to pattern setting enhancers (which contain binding sites for both transcriptional activators and repressors, Small and Arnosti 2020), developmental genes are regulated by the Polycomb (PcG) and Trithorax group genes (TrxG). Studies in Drosophila show that PcG and TrxG genes can impart a memory of the early pattern by setting the chromatin in an “ON” or “OFF” transcriptional state (reviewed in Kassis et al. 2017; Schuettengruber et al. 2017). We are interested in how chromatin environment influences the enhancer activity of developmental genes.

The Drosophila *engrailed* (*en*) gene encodes a homeodomain transcription factor whose best-known functions are in embryonic segmentation and specification of the posterior compartment in larval imaginal discs, precursors of the external structures of the adult (Garcia-Bellido and Santamaria, 1972; Morata and Lawrence, 1975; Kornberg 1981). En is expressed in the embryo in a series of stripes in the ectoderm, and subsets of cells in the central and peripheral nervous systems, hindgut, fat body, posterior spiracles, and head (Dinardo et al. 1985). Using a reporter gene in transgenic flies, we identified 20 embryonic enhancers spread over a 66kb region including DNA upstream, within, and downstream of the 4kb *en* transcription unit (Cheng et al 2014). However, we were not able to identify a fragment of DNA that drove expression of a reporter gene in the posterior compartment of imaginal discs in an *en*-like pattern. We speculated that, like the imaginal disc enhancers of the *Ultrabithorax* (*Ubx*) gene (Poux et al 1996; Pirrotta et al 1995; Müller and Bienz, 1991; Christen and Bienz 1994), the ‘ON-OFF’ state of the *en* imaginal disc enhancers is set by the embryonic expression pattern and remembered throughout development through epigenetic memory; without this epigenetic memory, the *en* imaginal disc enhancers could not regulate a reporter gene in the appropriate pattern.

*en* exists in a gene complex with *invected* (*inv*). *inv* encodes a closely related homeodomain protein that is largely co-regulated with *en* (Coleman et al. 1987; Gustavson et al. 1996). In the ‘OFF’ transcriptional state, H3K27me3, the repressive chromatin mark put on by the Polycomb protein complex PRC2, covers the entire *inv-en* domain, showing that *inv-en* is a target for Polycomb-mediated repression (Schwartz et al. 2006; De et al. 2016). Consistent with this, Polycomb group genes (PcG) are required to silence *inv-en* expression where they are not normally expressed in embryos and imaginal discs (Bustaria and Morata 1988; Moazed and O’Farrell 1992; Randsholt et al. 2000; Nekrasov et al. 2007). In our dissection of *inv-en* regulatory DNA we were surprised to find two fragments of DNA that predominantly caused reporter gene expression in the anterior but not the posterior compartment in the wing imaginal disc, the opposite of the En pattern (Cheng et al. 2014). We reasoned that the reporter gene was being expressed in the anterior compartment because Polycomb silencing was not established over the imaginal disc enhancer. But what was silencing reporter gene expression in the posterior compartment? Overexpression of En via an inducible transgene silences En expression in imaginal discs (Guillén et al. 1995; Tabata et al. 1995). We hypothesize that taking the *en* imaginal disc enhancers outside of the *inv-en* domain made them more susceptible to repression by En.

Here we study two imaginal disc enhancers for En using three approaches: 1) testing their activity in reporter constructs, 2) deleting them from *HAen79*, a 79kb transgene with HA-tagged En, that can rescue *inv-en* double mutants (De et al. 2019), and 3) deleting them from *invΔ33*, a chromosome that contains a 33kb deletion of *inv* DNA, creating a mimic at the endogenous *en* locus of the sequences present in *HAen79* (called *en80* in De et al. 2019). Our results suggest that the En protein directly represses its own expression through the imaginal disc enhancers and other sequences within the *inv-en* domain. Deletion of either imaginal disc enhancer from the *HAen79* transgene causes a loss-of-function *en* phenotype. In contrast, the same deletions do not cause phenotypes when deleted from the *invΔ33* endogenous locus. Adding an insulator to one end of the *HAen79* transgene increases the function of this transgene. Altogether our experiments show that the imaginal disc enhancers are epigenetically regulated and that their function depends in part on the chromatin environment of the endogenous *inv-en* domain.

## Results

The *inv* and *en* genes are contained within a 113kb domain flanked by the genes *E(Pc)* and *tou* (Fig. 1). *en* is required for both embryonic and adult development. In contrast, the *inv* gene is not required for viability or fertility in the laboratory (Gustavson et al. 1995). Inv and En are co-expressed in most cells in embryos and imaginal discs (Gustavson et al. 1995; Cheng et al. 2014), although some cells express Inv more strongly than En, and vice versa. For example, in wing imaginal discs, where Inv and En are co-expressed in cells in the posterior compartment of the disc, a group of cells in the hinge region express Inv at a much higher level (Fig. 1B, arrow). In addition, the amount of Inv and En is not uniform in the posterior compartment of the wing disc. We point this out to raise the possibility that En and Inv might regulate both their own and each other’s expression levels.

**Fig. 1.**
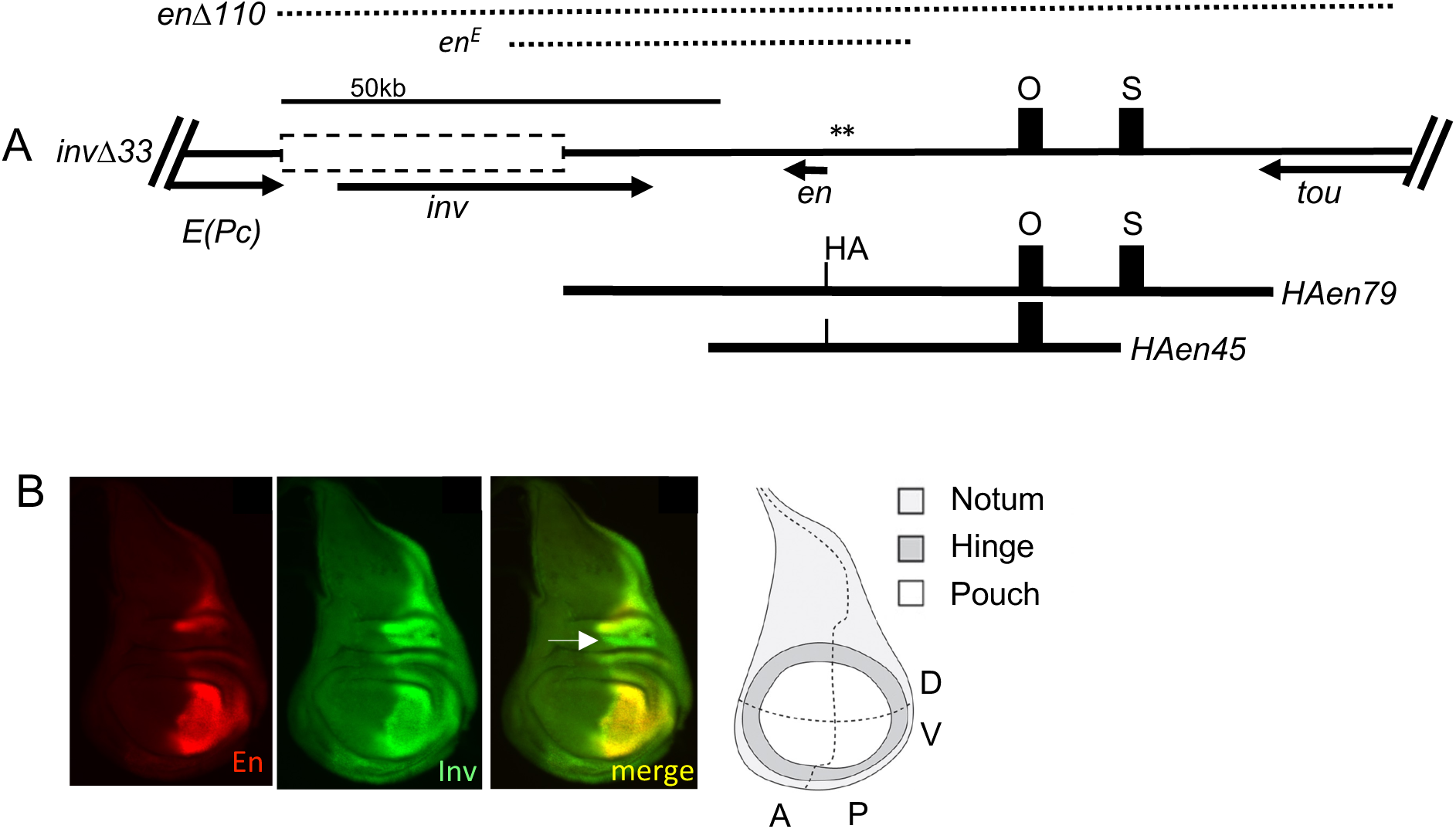
Map of *invΔ33*, transgenes, PREs and imaginal disc enhancers. **(A)** Diagram of the *inv- en* region of the genome with flanking genes. The 33kb region deleted on the *invΔ33* chromosome is outlined in a dashed box (*en80* in De et al. 2019). The black boxes labeled ‘O’ and ‘S’ are the location of the enhancers studied in this paper. Asterisks (**) denote the locations of the constitutive (or “major” De et al. 2016) *en* PREs. The *inv* PREs are not shown because they have been deleted in *invΔ33*. The arrows denote the direction and extent of the transcription units. The DNA deleted in *enΔ110* and *en^E^* is shown by dotted lines. Bottom, the DNA present in the two θ>C31 transgenes used in this study. In these transgenes, En is labeled on the N-terminus with a single HA-tag (Cheng et al. 2014). (B) Expression pattern of En and Inv in a wild-type wing disc. White arrow points to a region of the disc where Inv protein is higher than En. A fate map of a wing imaginal disc is shown on the right. A-anterior, P- posterior, D-dorsal, V-ventral. (Diagram from Simon and Guerrero 2015).

### Fragments O and S are imaginal disc enhancers

The locations of two fragments of DNA, ‘O’ and ‘S’, that gave expression in imaginal discs in a reporter construct are shown in Fig 1. To test the hypothesis that the ‘ON-OFF’ state of these enhancers could be set at the embryonic stage, we cloned them in a vector that gives striped expression throughout most of embryogenesis but not in imaginal discs (construct H, Cheng et al. 2014, Fig. S1). En embryonic stripes are driven by different enhancers as development proceeds (DiNardo et al. 1988; Heemskerk et al. 1991; Cheng et al. 2014). Fragment O is 3.9kb and, in addition to the imaginal disc enhancer, contains some stripe enhancers, including for stage 11, the only stage missing from the stripe enhancers present in construct H. For ‘S’, we used a 2.8kb fragment, considerably smaller than the 6.7kb fragment we previously studied (Cheng et al. 2014). The coordinates of this fragment were set by an overlap of our original ‘S’ fragment and an imaginal disc enhancer identified in a screen of genomic fragments for cis-regulatory activity in imaginal discs (line GMR94D09, Jory et al. 2012). There are no embryonic enhancers present in this 2.8kb fragment ‘S’.

We reasoned that the stripe enhancers present in construct H might ‘set’ the enhancers present in fragments O or S in the “ON” or “OFF” state, leading to expression of ß-galactosidase (ßgal) in the posterior compartment of wing discs. For ‘S’, this did not occur. The expression of ßgal in three independent insertion lines was stronger in the anterior than the posterior compartment of the wing disc (Fig. S1). This was most dramatic in one line where the expression of ßgal was nearly missing where En was expressed. These data are consistent with the model that ‘S’ contains an enhancer for the entire wing disc but is repressed in cells expressing En. ßgal expression from the O construct was quite variable. In one line, ßgal was off in the anterior compartment, like En, but only partially on in the posterior compartment (Fig. S1). In another, ßgal was expressed in the posterior compartment, and mostly silenced in the anterior, and in another, anterior expression was stronger than posterior, similar to expression driven by fragment ‘S’. These data suggest that enhancer ‘O’ might also be repressed by En, and that the extra stripe enhancers present in the ‘O’ fragment (including stage 11) might be able to set the “ON” and “OFF” states better than fragment ‘S’ that lacks stripe enhancers. For both constructs, the variability in expression patterns seen between lines is consistent with the view that these enhancers are regulated by the chromatin environment of the transgene insertion site. Finally, although in this paper we describe the activity of fragments ‘S’ and ‘O’ in wing discs, both these enhancers also drive expression in all other discs examined (haltere, leg, and eye-antennal discs, Fig. S2).

We cloned the ‘S’ fragment into a different vector used to detect enhancer activity and dissected it into two smaller fragments, SS2 and SS1 (Fig. 2). Chromatin-immunoprecipitation followed by sequencing (ChIP-seq) in 3^rd^ instar larval brains and discs show the location of the Polycomb proteins Pho, Ph, and the En protein and the H3K27me3 chromatin mark over the DNA present in the *invΔ33* allele (Fig. 2A). Normally, En expression is silenced by Polycomb proteins in the anterior compartment in discs (Bustaria and Morata 1988; Randsholt et al. 2000; Nekrasov et al. 2007), consistent with H3K27me3 covering this region of the chromosome in this mixed cell population. ‘S’, ‘SS2’, and ‘SS1’ were cloned in front of the GAL4 reporter gene (Fig. 2C, D) and integrated into two different insertion sites: attP40 and attP2. At both chromosomal locations, ‘S’ and ‘SS2’ gave nearly identical expression patterns, in the anterior compartment, although expression from ‘SS2’ was weaker. There is an En ChIP-seq peak directly over the SS2 fragment (Fig. 2B). These data suggest that En may directly repress the expression of the transgene in the posterior compartment. ‘SS1’ has no enhancer activity in wing discs (Fig. 2D).

**Fig. 2.**
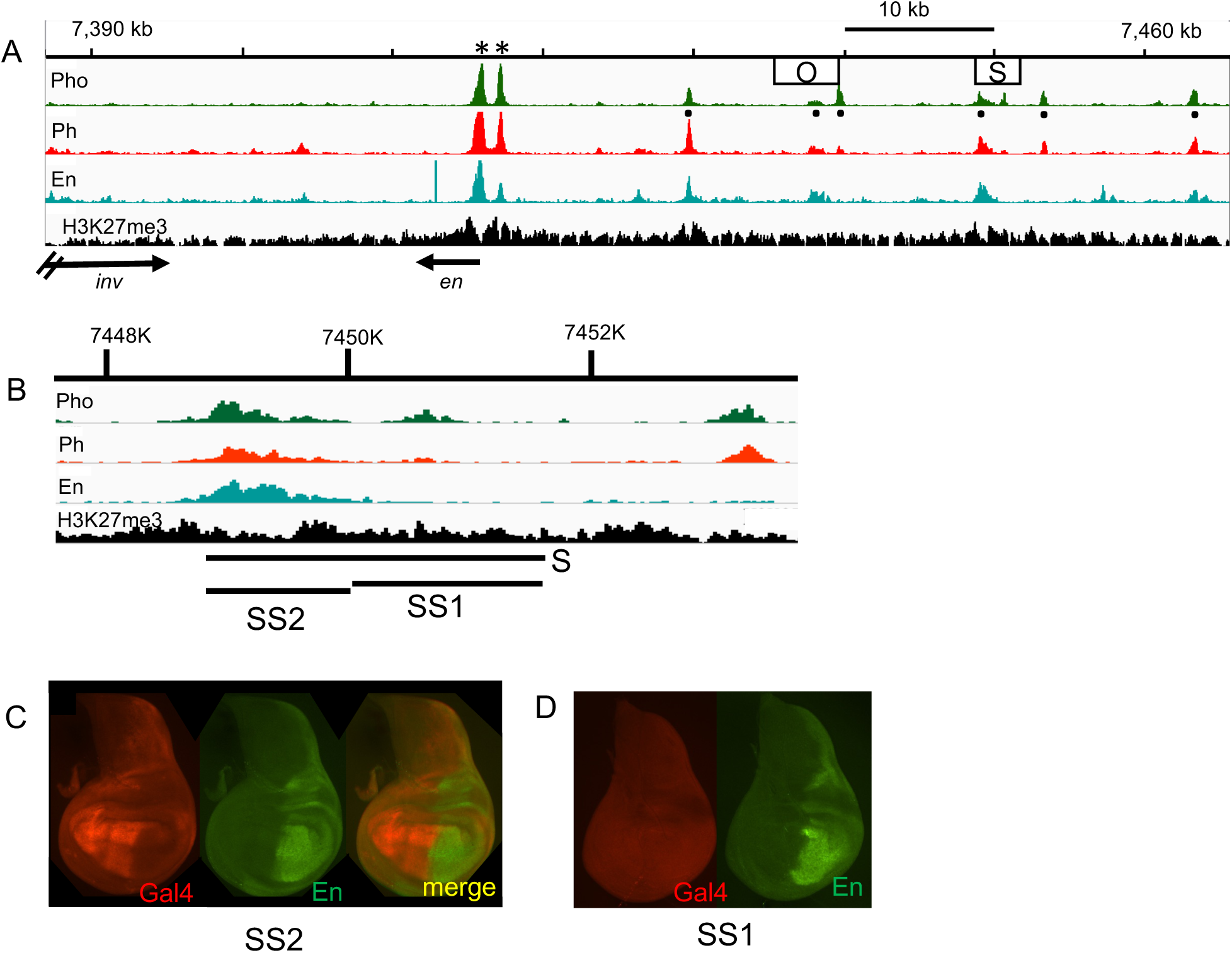
Activity of enhancers ‘O’ and ‘S’ in a reporter construct. (A) ChiP-seq data on 3^rd^ instar larval brains and discs for Pho, Ph, En, and H3K27me3 over the genomic region present in *en80* (Coordinates chr2R, version dm5; GSE76892, Lorberbaum et al. 2016). Asterisks indicate the position of the constitutive (aka major) *en* PREs. “Minor” or tissue specific PREs are marked by black dots below the Pho ChIP-seq peaks. These could be dual function elements that serve as enhancers or silencers dependent on the context (Erceg et al. 2017). Locations of the O and S fragments are shown as boxes. (B) Expanded view of the region around fragment S showing the locations of the SS2 and SS1 fragments. (C,D) Gal4 (red) expression in wing discs from transgenic flies containing SS1 or SS2 cloned in front of Gal4 (in *pBPGUw*, Pfeiffer et al. 2008). En (green) is shown for comparison. These transgenes were inserted at attP40. Similar results were obtained with the same transgenes inserted at attP2.

We next turned our attention to two large HA-en transgenes, one 45kb and one 79kb (see Fig. 1A). We previously showed that *HAen45* can rescue *en* point mutants, but not double mutants of *inv* and *en* (Cheng et al. 2014). The *HAen79* transgene completely rescues *inv-en* double mutants, including *enΔ110* that deletes the most of the *inv-en* domain (De et al. 2019). Despite this, the expression of HA-en from the transgenes is not normal in wing discs in a wildtype background (Fig. 3). In *HAen45*, HA-en is nearly absent in the pouch region of the wing disc at three different attP insertion sites, and variegated at the other (Fig. 3A, E). Expression of HA-en in the *HAen45* wing pouch is partially restored in the presence of the *en^B86^* mutation, a deletion of 53bp in the coding region of En, that produces no detectable En protein (Gustavson et al 1995; Fig. 3B). These data suggest that the repression of *HAen45* is mediated by the En protein itself and not via an interaction of the transgene with the endogenous *inv-en* gene, as has been seen at the Drosophila *spineless* gene (Viets et al., 2019). *HAen79* is expressed better than *HAen45*, but there are still regions of the wing disc where it is not expressed (Fig. 3C, arrow). The size of the repressed region is variable from disc to disc, and the expression is variegated. This variegation is dramatic when *HAen79* is inserted at attP3 (Fig. 3E). We suggest that this variegated expression is the result of unstable gene expression, a competition between the “ON” and “OFF’ transcription states set by epigenetic marks.

**Fig. 3.**
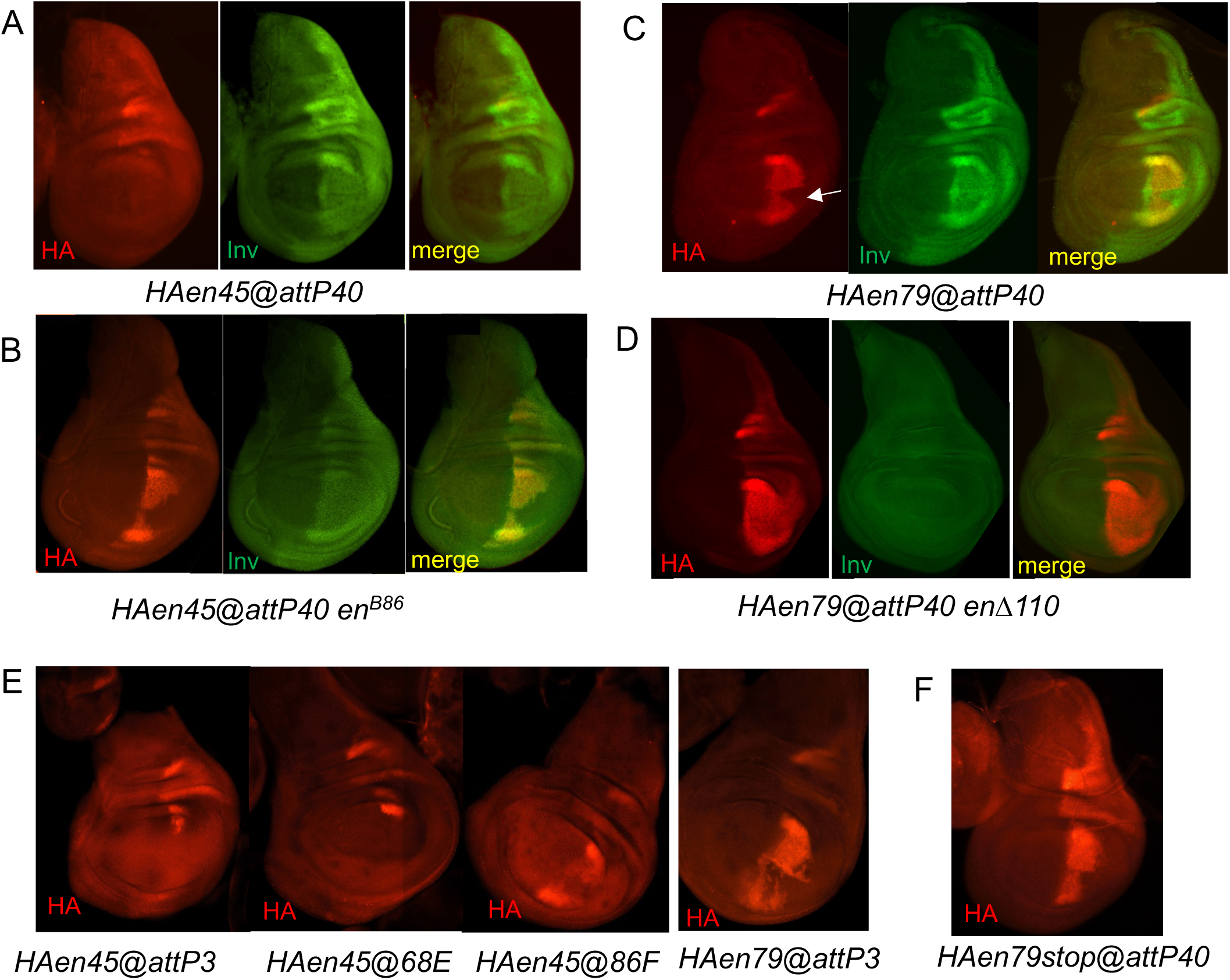
HA-en expression from transgenes is repressed by wildtype levels of En. (A,B) Expression of HA-en in wing discs from *HAen45* in a wildtype background (A) and on a chromosome with *en^B86^* (B). *en^B86^* makes no En protein but expresses Inv in a wild-type pattern. ((B) is reprinted from Cheng et al. 2014) (C, D) Expression of *HAen79@attP40* in a wildtype background (C) and when on the same chromosome as *enΔ110* (D). *enΔ110* deletes the entire *inv-en* domain so Inv is not present. The white arrow points to a place where HA-en is not present. (E) HA-en expression from the *HAen45* and *HAen79* transgenes at different insertion sites. (F) *HAen79stop* expresses a non-functional HA-en protein. All discs are homozygous for the indicated genotype.

We also asked whether the HA-en made by the transgene is necessary for its variegated expression. We made *HAen79-stop* that contains a stop codon in En and produces a non-functional En protein (De et al. 2019). Like *HAen79@attP40*, *HAen79-stop@attP40* is expressed in only part of the wing pouch (Fig. 3F) and its expression is variegated. We conclude that the *HAen79* transgene is repressed by En expressed from the wildtype *inv-en* domain and that the HA-en protein contributes very little to this repression.

We next deleted fragments O, S, or SS2 from the *HAen79* transgene, inserted them at *attP40*, and compared the expression of HA-en in wing imaginal discs, both with and without wildtype Inv and En (Fig. 4). Deleting fragment O decreases expression of HA-en in a wildtype background, especially in the ventral region of the wing pouch (Fig. 4A, white arrow). The O fragment contains an embryonic stripe enhancer for the ventral part of the embryo at stage 12 of development (Cheng et al. 2014). Our data suggest either that ‘O’ also contains an enhancer for the ventral wing disc, or that epigenetic memory is impaired when this embryonic enhancer is removed. Deleting fragments ‘S’ or ‘SS2’ from *HAen79* gave essentially the same result in wing discs (Fig. 4A). In a wild-type background, expression is only observed in a line at the anterior-posterior (A-P) boundary. There are three different enhancers for this A-P boundary line present in *HAen79*, and they are not within fragments ‘O’ or ‘S’ (Cheng et al. 2014). Removal of the *inv-en* domain leads to almost wild-type expression from *HAen79ΔO* and more, but still variegated, expression of *HAen79ΔS* or *HAen79ΔSS2* (Fig. 4A). The wing phenotypes of *HAen79ΔO*, *ΔS* or *ΔSS2 enΔ110* are consistent with these expression patterns (Fig. 5A). Minor wing vein defects are seen in *HAen79ΔO enΔ110* wings, and more severe phenotypes are seen in *HAen79ΔS enΔ110* and *HAen79 ΔSS2enΔ110* wings, including the presence of anterior-like bristles on the posterior wing margin, indicating a posterior-to-anterior transformation (Fig. 5A). Altogether these data show that fragments ‘O’, ‘S’, and ‘SS2’ contain wing disc enhancers.

**Fig. 4.**
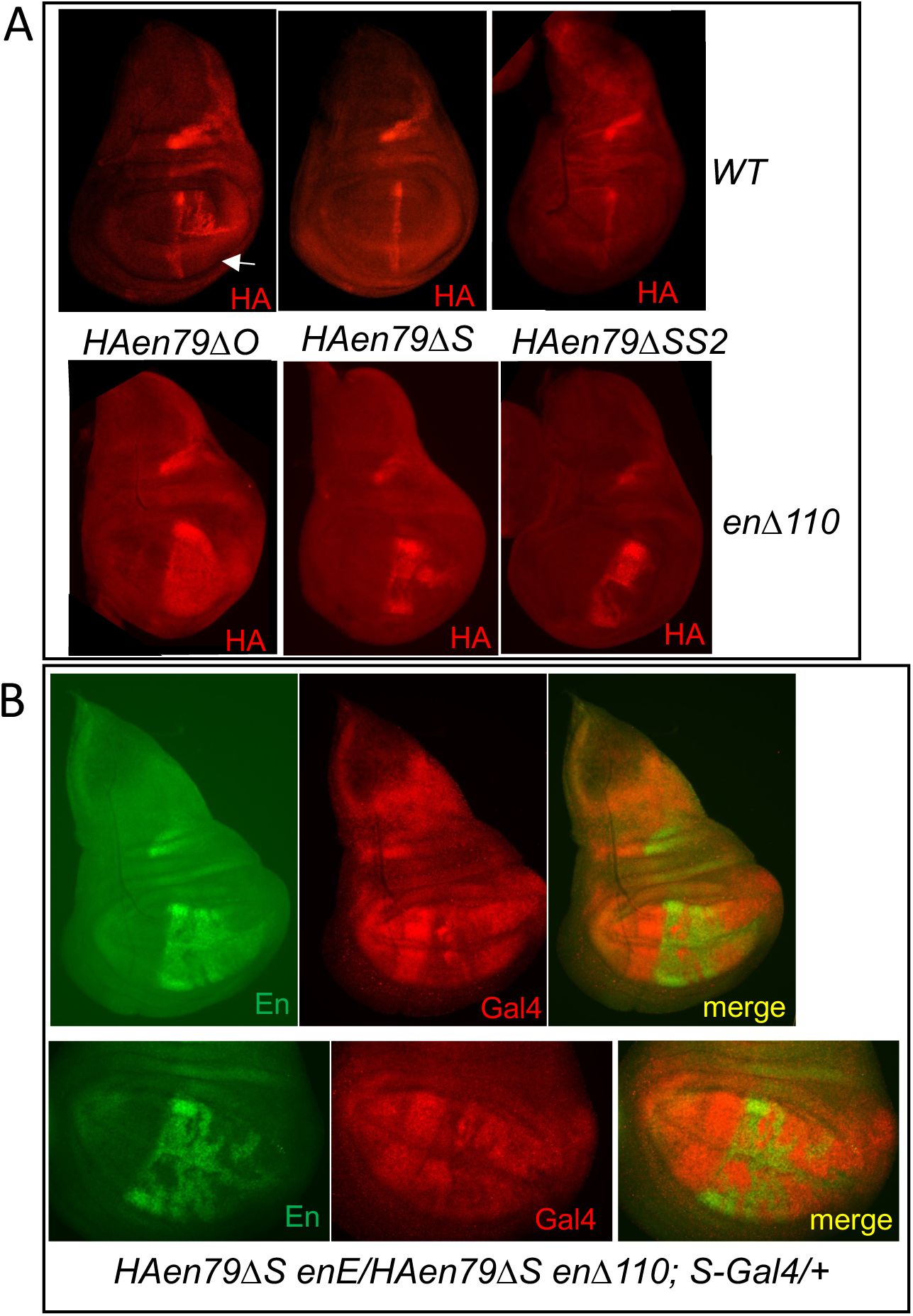
En represses expression of fragment O and S containing transgenes. (A) Top row: HA expression from *HAen79ΔO*, *HAen79ΔS*, and *HAen79ΔSS2* (all inserted at attP40) on a wildtype chromosome. Bottom: The same transgenes on an *enΔ110* chromosome. All discs are homozygous for the indicated genotype. Arrow points to the ventral portion of the wing disc. (B) En (green) and Gal4 (red) in two different wing imaginal discs of the genotypes *HAen79ΔS enE/HAen79ΔS enΔ110; S-gal4/+*. The bottom row shows a close-up of the pouch region of a different wing disc. *HAen79ΔS* is the only source of En in this genotype and is expressed in a variegated manner in the wing pouch. S-gal 4 is expressed in the wing pouch predominantly in cells that do not express En.

**Fig. 5.**
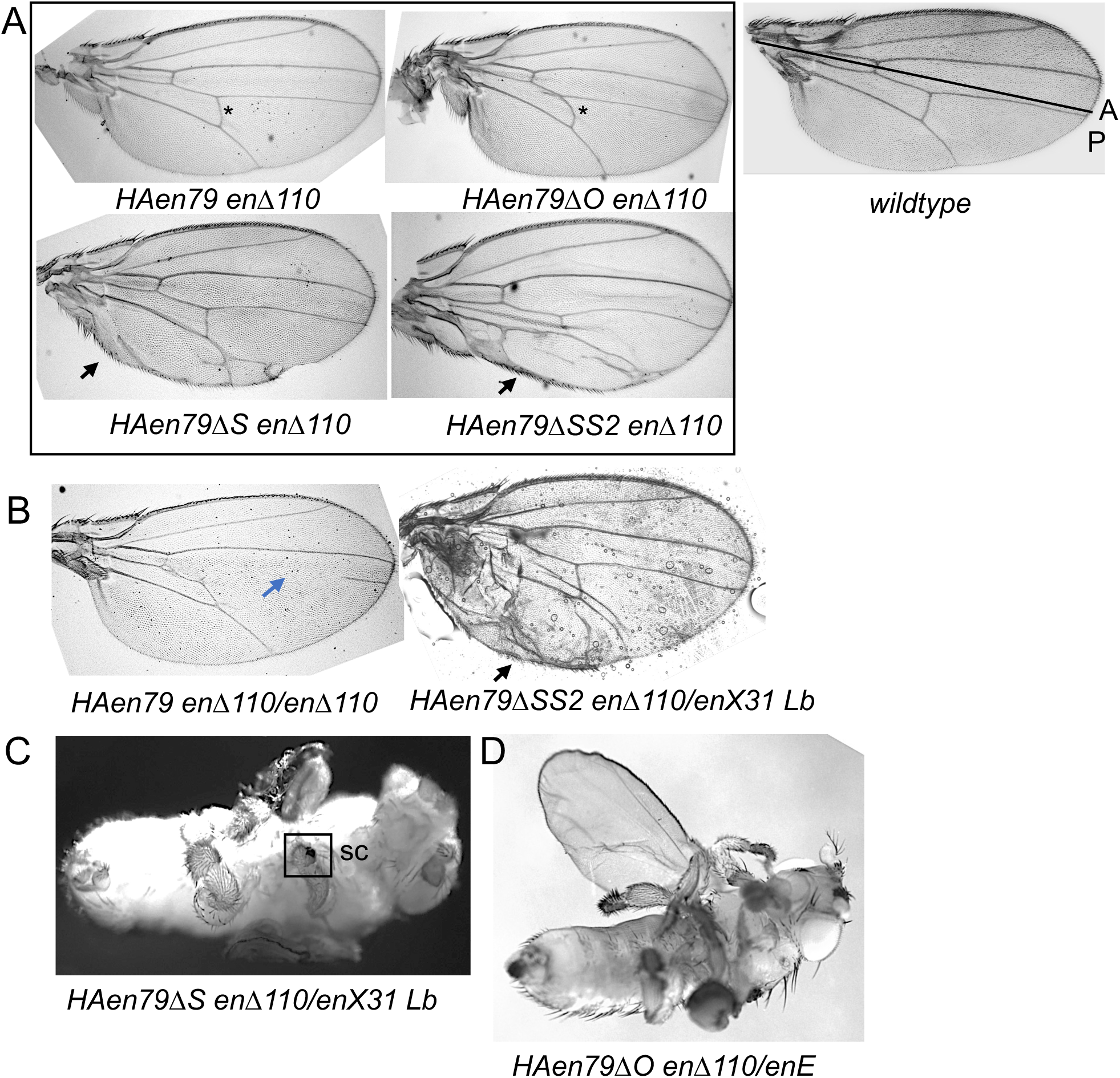
*HAen79* transgenes are haploinsufficient. (A) Wings of flies homozygous for the indicated genotypes. Asterisk marks the crossvein where minor defects are seen. Arrows point to the presence of anterior margin bristles on the posterior wing margin. A wildtype wing is shown with a line separating the A (anterior) and P (posterior) compartments. (B) Wings from flies of the indicated genotypes. The blue arrow points to the missing wing vein. The black arrow points to anterior bristles on the posterior wing margin. (C) Ventral side of pharate adults of the genotypes shown. On the left, two legs are visible; the middle leg has been removed. The boxed region is the sex comb teeth (sc). The distal part of this leg is missing and the proximal part is misshapen. This leg phenotype was extreme but seen in many *HAen79ΔS enΔ110* or *HAen79ΔO enΔ110/enX31* pharates. (D) This pharate has an extended wing missing many veins and less severe leg defects.

### A strong correlation between En protein and repression of the ‘S’ enhancer

We used the variegated expression of HA-en from *HAen79ΔS* as a tool to address the correlation between En expression and repression of the ‘S’ enhancer. We constructed a genotype *HAen79ΔS enΔ110/ HAen79ΔS en^E^; S-Gal4@attP2/+* and examined En and Gal4 distribution in wing discs. *en^E^* is a 41kb deletion that removes En and produces a truncated Inv protein that lacks the homeodomain (Fig. 1A). In this background, the only source of En is from the *HAen79ΔS* transgene. Strikingly, in the posterior compartment of the wing pouch, *S-Gal4* is repressed in the cells where En is expressed (Fig. 4B). These data, along with ChIP data that show En binding to ‘S’, support the hypothesis that En can directly repress the S-enhancer in the wing pouch.

### One copy of the *HAen79* transgene is not enough

Flies survive well with one copy of the *inv-en* domain and have no known phenotypes. That is not the case with the *HAen79* transgenes. The phenotypes of *HAen79* and its derivatives as homo- and heterozygotes are shown in Fig. 5. While homozygous *HAen79 enΔ110* flies survive with only minor wing defects, flies with only one copy of *HAen79* in a homozygous *enΔ110* background have wing defects and survive poorly (Fig. 5B and Table 1). Some *HAen79 enΔ110/enΔ110* flies hatch and then stick to the sides of the vial or fall in the food and die, suggesting a defect in nervous system development. In contrast, *HAen79ΔO* or *HAen79ΔS enΔ110/inv-en* deletions die as pharate adults with severe leg defects, and wings that are usually not expanded (Fig. 5C). A rare *HAenΔO enΔ110/en^E^* fly with an expanded wing showed a lack of wing veins throughout most of the wing and deformed legs (Fig. 5D). *HAen79ΔSS2 enΔ110/inv-en* deletion flies survive with wing defects similar to those seen in *HAen79ΔSS2 enΔ110* homozygotes and have no leg defects. These data suggest that ΔS takes out more regulatory sequences than does ΔSS2, consistent with the higher expression level of S-gal4 than SS2-gal4. Thus, although the *HAen79* transgene can rescue *inv-en* double mutants, it is a not equivalent to a wildtype *en* locus. Deleting either the ‘O’ or ‘S’ fragment from the *HAen79* transgene leads to leg defects when these transgenes are the only source of En (Fig. 5C, D). This provides further evidence that these fragments are also enhancers in leg discs.

**Table 1:** Viability and phenotype of flies with a single copy of the *en* gene.

| Genotype of fathers | Genotype of mothers |  |  |  |  |  |  |  |  |
| --- | --- | --- | --- | --- | --- | --- | --- | --- | --- |
|  | *CyO* / *enΔ110* |  |  | *CyO* / *en<sup>E</sup>* |  |  | *CyO* / *enX31* |  |  |
|  | Progeny |  |  | Progeny |  |  | Progeny |  |  |
|  | *CyO* | *EnΔ110* |  | *CyO* | *en<sup>E</sup>* |  | *CyO* | *enX31* |  |
|  | # | # | Phenotype | # | # | Phenotype | # | # | Phenotype |
| *invΔ33* | 68 | 77 | + | 51 | 34 | + | 63 | 36 | Wings out |
| *invΔ33ΔO* | 46 | 55 | Wings out | 48 | 46 | Wings out | 84 | 46 | Wings out |
| *invΔ33ΔS* | 37 | 41 | Wings out | 68 | 77 | + | 106 | 44 | Wings out |
| *invΔ33ΔSS2* | 46 | 47 | Wings out | 41 | 42 | + | 99 | 45 | Wings out |
| *invΔ33ΔOΔSS2* | 94 | 79 | Wings out** | 73 | 96 | Wings out** | ND | ND | ND |
| *HAinvΔ33DSS2* | 44 | 46 | Wings out | 54 | 34 | + | 99 | 45 | Wings out |
| *HAen* | 41 | 59 | + | 48 | 51 | + | 68 | 35 | + |
| *HAen79 enΔ110* | 146 | 9* | Wings out** | 136 | 2* | Wings out ** | 80 | 2* | Wings out** |
| *HAen79GyB enΔ110* | 151 | 86 | Wings out | 110 | 98 | Wings out | 75 | 41 | Wings out |
| *HAen79GyMW enΔ110* | 79 | 98 | Wings out | 77 | 76 | Wings out | 93 | 41 | Wings out |

+ Wildtype, \* A few flies hatch and either fall in the food and die or are stuck to the pupal case or sides of vial.
\*\* Wing vein defects in posterior compartment (a loss of function phenotype).

### En expression from *invΔ33* is not sensitive to the loss of the ‘O’ or ‘SS2’ enhancers

*invΔ33* was created as a mimic of the *HAen79* transgene at the endogenous locus (called *en80* in De et al. 2019, Fig. 1). At *invΔ33*, in addition to the 79kb present in *HAen7*9, there is 1kb of DNA just downstream of the *E(Pc)* transcription stop site. *E(Pc)* and *tou* transcription form the boundaries of the *inv-en* domain (De et al. 2020). We left 1kb downstream of *E(Pc)* because we did not want to risk interfering with *E(Pc)* transcription termination. We used *CRISPR/Cas9* to delete either fragment O or SS2 from *invΔ33* and saw no difference in En expression in wing discs (Fig. 6). We next tested whether *invΔ33*, *invΔ33ΔO* or *invΔ33ΔSS2* were sufficient as single copies by crossing them to three *inv-en* deletion mutants (Table 1). All three lines survive well over all the deletion mutants; none have wing vein defects or leg defects. Some flies hold their wings out, a phenotype associated with loss of *inv* (De et al. 2016). We wondered whether adding an HA-tag to En would impair its function. We HA-tagged En on both a wildtype chromosome and on *invΔ33ΔSS2*, making *HAinvΔ33ΔSS2*. We found no evidence that the HA-tag compromised En function, as *HAinvΔ33ΔSS2* flies survive well as heterozygotes and do not have wing vein defects (Table 1). In summary, these data show that the endogenous *invΔ33* domain is resilient to the loss of a single imaginal disc enhancer.

**Fig. 6.**
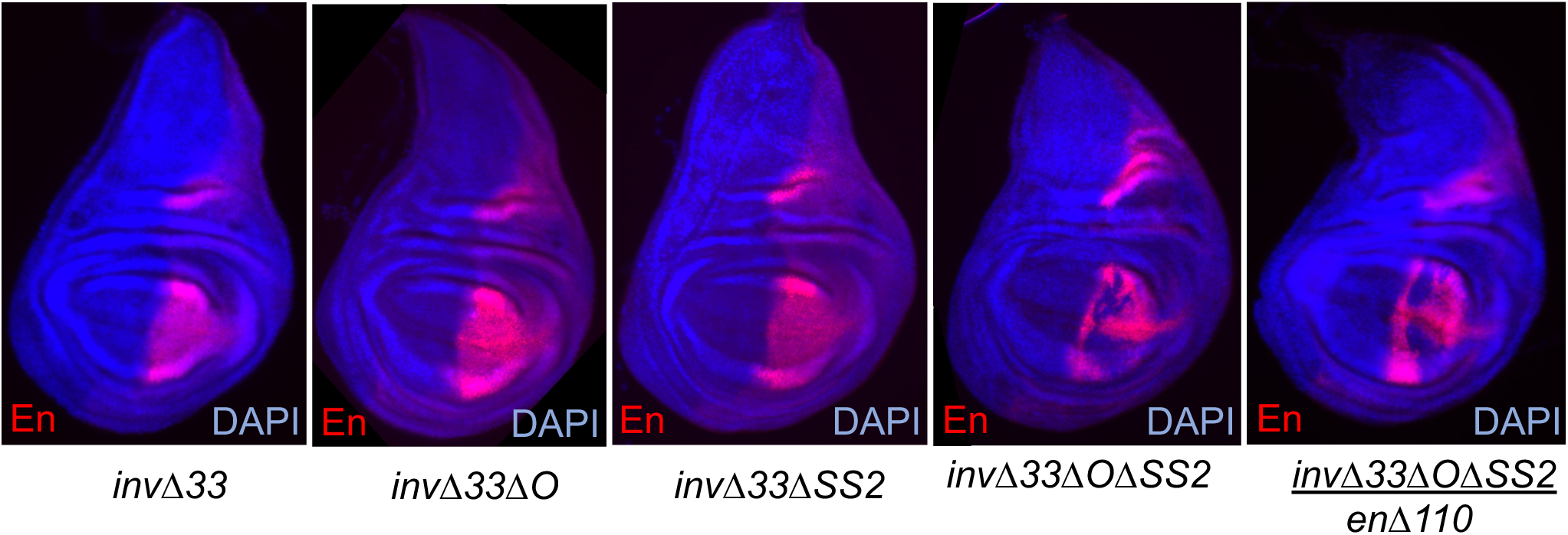
Deletions of disc enhancers from *invΔ33* reveals the stability of the endogenous locus. En in *invΔ33*, *invΔ33ΔO* and *invΔ33ΔSS2* wing imaginal discs looks like WT. En in *invΔ33ΔOΔSS2* homozygous and *invΔ33ΔOΔSS2/enΔ110* wing discs is variegated.

We next deleted both fragments O and SS2 and found that En expression is variegated both in *invΔ33ΔOΔSS2* homozygotes and *invΔ33ΔOΔSS2/enΔ110* wing discs (Fig. 6). This variegated expression is consistent with the hypothesis that these enhancers are regulated by chromatin modifications*. invΔ33ΔOΔSS2* survive well as heterozygotes (Table 1). Consistent with the variegated expression patterns, some wings have vein defects, whereas others do not.

We hypothesized that the endogenous locus was more stable to enhancer deletions than the HA-en transgene because it has boundaries that stabilize the chromatin state of the locus. For example, H3K27me3, the Polycomb chromatin mark, stops at the 3’ ends of the *E(Pc)* and *tou* genes at the endogenous locus (De et al. 2020). On the other hand, the transgene does not have boundaries, and the H3K27me3 spreads from the *en* DNA in both directions many kilobases, stopping at actively transcribed genes (De et al. 2019). We hypothesize that this destabilizes the transgene in both the “ON” and “OFF” transcriptional states, making it less stable and more sensitive to the loss of enhancers.

### Adding a gypsy element boundary to *HAen79* improves its function

The *HAen79* transgene also contains a mini-*white* (*mw*) gene as a marker to detect integration events into an *attP* landing site (Venken et al. 2006). A mini-*yellow* (*y*) gene is present at the landing site and is located downstream of the *en* transcription unit after integration of the *HAen79* into the genome (Fig. 7A). We used *CRISPR/Cas9* to insert gypsy boundary elements at the ends of the 79kb HA-*en* DNA. Gypsy boundaries can block both the action of enhancers and the spreading of the H3K27me3 Polycomb mark (Mallin et al. 1998; Scott et al. 1999; Kahn et al. 2006). Two lines were created, one with gypsy on the *mw* side, *HAen79GyMW*, and the other with gypsy on both sides, *HAen79GyB*. In contrast to flies with just one copy of *en* from the *HAen79enΔ110* chromosome, both *HAen79GyMW enΔ110*, and *HAen79GyB enΔ110* flies survive well over deficiencies for *inv* and *en* (Table 1) and have no defects in wing veins. Unlike the original *HAen79* line that has variegated expression in the wing pouch in the presence of a wildtype *inv-en* domain, both *HAen79GyMW* and *HAen79GyB* have HA-en expression in an En-like manner in the wing pouch (Fig. 7B). Unexpectedly, Inv expression is now variegated in these discs. This finding suggests that the HA-en expressed from the transgene is now strong enough to repress the expression of Inv at the endogenous locus.

**Fig. 7.**
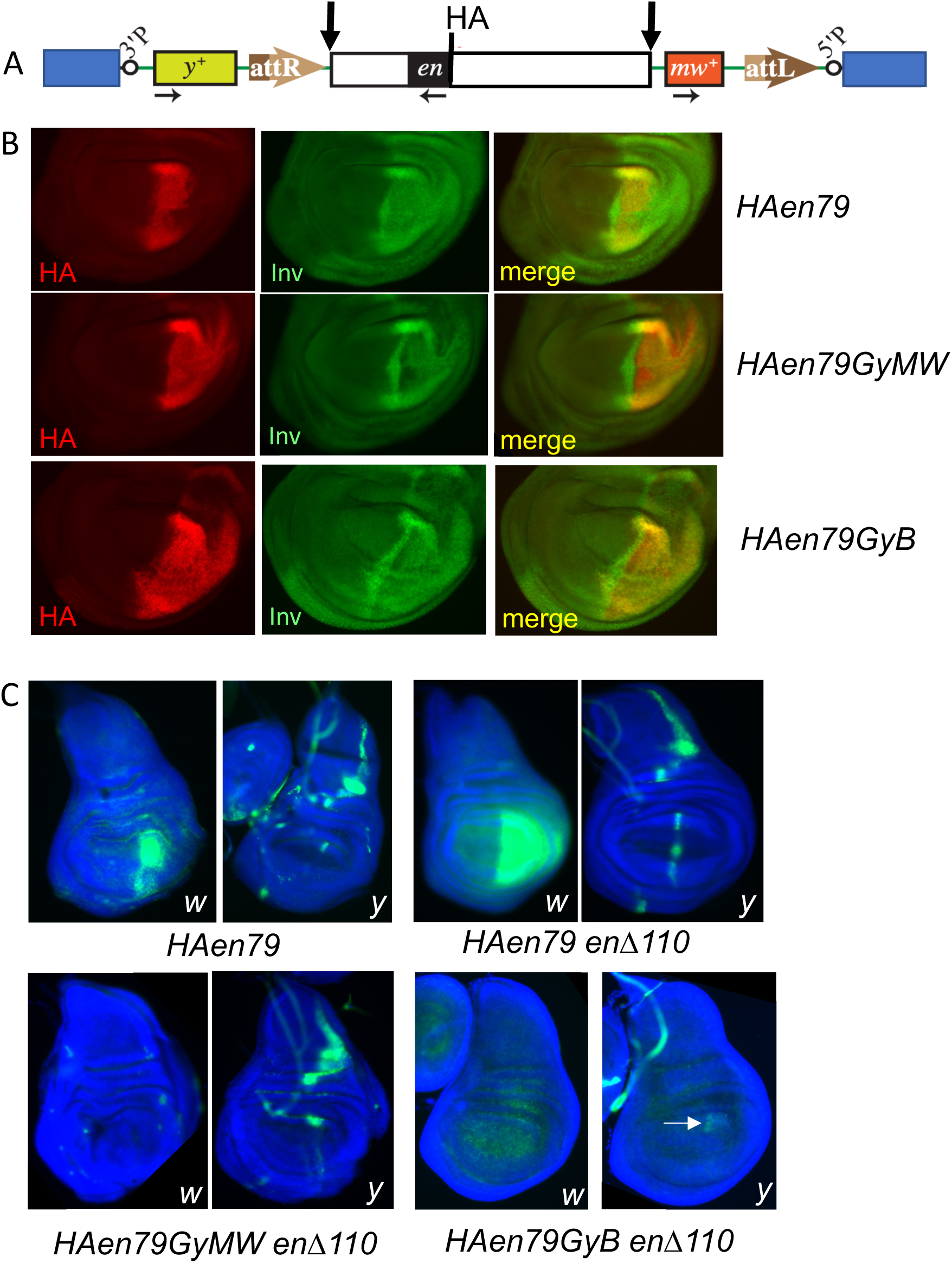
Flanking *HAen79* by gypsy elements stabilizes HA-en expression and restricts expression of flanking genes. (A) Diagram of the *HAen79* transgene (not to scale). The blue boxes are flanking genomic DNA. The *en* transcript is marked by a black box, the direction of transcription is shown by horizontal arrows. Vertical arrows indicate the gypsy boundary insertion sites. (B) HA and Inv expression in wing disc pouch from the genotypes indicated (on a wildtype chromosome). Note the expanded HA expression and variegated Inv expression in *HAen79GyMW* and *HAen79GyB* compared to *HAen79*. (C) RNA in situ hybridization. Note that both the mini-*white* (*w*) and *yellow* (*y*) genes are transcribed at higher levels in the wing pouch when the endogenous *inv-en* domain is deleted (*HAen79 enΔ110*). Adding a gypsy element on the mini-*white* site completely blocks *w* expression. A small amount of *y* expression remains (white arrow) in *HAen79GyB enΔ110*.

We next looked at expression of *white* (*w*) and *yellow* (*y*) RNA, the two genes that flank the 79kb HA-en transgene. In a wildtype background, *w* is expressed in a variegated manner in the wing pouch, like HA-en (Fig. 7C). *y* is expressed mainly in the notum part of the wing disc. When the *HAen79* transgene is the only *en* or *inv* gene in the genome (*HAen79 enΔ110*), *w* is expressed in the entire wing pouch, while *y* is expressed in a line at the anterior-posterior boundary in addition to the notum (Fig. 7C). Inserting a gypsy element between the *en* DNA and mini-*white* causes a complete loss of *w* expression, showing that gypsy is sufficient to block *en* enhancers from stimulating *w* expression. *y* is expressed in the notum and weakly at the A/P boundary in the wing pouch in *HAen79GyMW enΔ110* wing discs. Inserting gypsy elements on both sides blocks *w* expression, and almost completely blocks *y* expression (Fig. 7C).

We were surprised that the activity of *HAen79GyMW* is comparable to *HAen79GyB* for both viability as a heterozygote and HA-en expression in a wild-type background. However, perhaps this makes sense. Our previous data strongly suggested that all of the wing disc enhancers for expression in the posterior compartment are located upstream of the *en* transcription unit (Cheng et al. 2014). In addition, all of the Polycomb response elements (PREs), which also activate the expression of *HAen79* (De et al. 2019), are located upstream. Our data show that without the boundary between *en* and mini-*white*, the chromatin-regulated imaginal disc enhancers can activate the flanking *mw* gene. We suggest that this makes the overall stability of the “ON’ state weaker and subject to repression by the endogenous Inv/En proteins. This may also explain why loss of one enhancer from the transgene causes a phenotype not seen with a similar deletion in the *invΔ33* endogenous locus (see Discussion).

Finally, we examined the stability of Polycomb repression of the *HAen79* transgenes with and without the gypsy boundaries. We previously showed that either removing the *en* PREs from *HAen79* or putting the *HAen79* transgene in a *ph-p^410^* background (which reduces the amount of the PcG protein Polyhomeotic) leads to flies with disrupted abdomens (De et al. 2019). Adding a gypsy element to either one or both sides of *HAen79* reduces the abdominal phenotype seen in *ph-p^410^* flies (Fig. 8). The exact phenotype was variable and similar in both *ph-p^410^; HAen79GyMW* and *ph-p^410^; HAen79GyB* flies. The equivalent phenotypes obtained by flanking only one or both sides of *HAen79* with gypsy boundaries was surprising to us because H3K27me3 spreads in both directions past the *HAen79* ends into flanking DNA (De et al. 2019). We suggest that the enhancers necessary for the abdominal phenotype are located upstream of the *en* promoter where both the constitutive and tissue-specific PREs are present (De et al. 2016), creating a stable H3K27me3 domain in the upstream DNA. One final point, the abdominal phenotype of *invΔ33* flies in *ph-p^410^*is nearly wild-type. Removing the *en* PREs from *invΔ33* causes it to behave much differently in a *ph-p^410^*background: *ph-p^410^; invΔ33Δ1.5* flies have severely disrupted abdomens. These data show that the *en* PREs play an important role in stabilizing the *invΔ33* Polycomb domain.

**Fig. 8.**
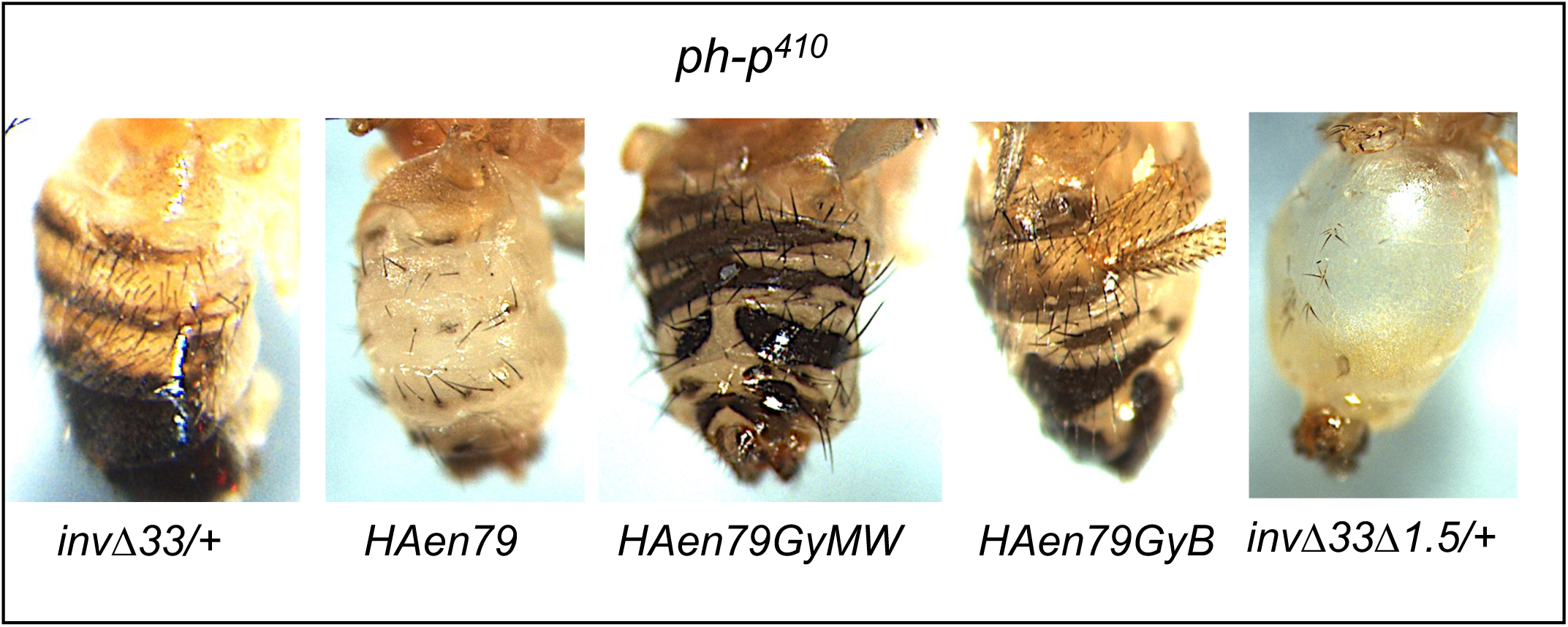
Adding gypsy elements to HAen79 improves the phenotype of the abdomen in *ph-p^410^* mutant males. *invΔ33*, *HAen79*, *HAen79GyB, invΔ33enΔ1.5* ventral-lateral views; *HAen79GyMW* ventral view. Part of a leg is also evident in *HAen79GyB*. Only one copy of the transgene is present in these genotypes. *invΔ33* and *invΔ33enΔ1.5* are heterozygous with a wildtype chromosome. Adding gypsy on one or both sides of *HAen79* led to similar phenotypes in *ph-p^410^* that were more like wildtype. Removing a 1.5kb fragment including both constitutive *en* PREs (*invΔ33enΔ1.5,* called *en80Δ1.5* in De et al. 2019) leads to flies with a very disrupted abdomen in the *ph-p^410^* mutant background.

## Discussion

Here we have shown that the activity of two *en* imaginal disc enhancers is dependent on the chromosomal context. En is normally expressed in the posterior compartment in imaginal discs. When cloned into small transgenes, these enhancers induce expression mainly in the anterior compartment. En binds to both these enhancers and may act directly to repress their expression in transgenes. Deleting either enhancer from a 79kb-HA-en transgene caused defects in expression of HA-en in the posterior compartment and decreased its ability to function as the sole source of En. The same deletions in the endogenous *invΔ33* locus did not cause any obvious phenotypes. Inserting a gypsy boundary between the upstream En DNA and the mini-*white* gene in *HAen79* increases its activity, allowing it to rescue as a single copy, and increasing the stability of its expression when combined with a mutation in the PcG gene *ph*. Adding an additional gypsy boundary between the downstream *en* DNA and the mini-*yellow* gene did not increase the activity or stability of the transgene to a *ph* mutation further. This was surprising to us and suggests that, in this case, looping of boundary elements may not be required for formation of a stable epigenetic state. In both *invΔ33* and *HAen79*, the *inv* PREs are missing, while the enhancers for expression in the posterior compartment of the wing disc, the precursors of the adult abdomen, and the constitutive and tissue-specific PREs are all located upstream of the *en* transcription unit (to the right of *en* in Fig. 1 map). We therefore propose that by inserting gypsy to generate *HAen79GyMW*, thereby blocking the spreading of H3K27me3 upstream and blocking the enhancers from acting in the direction of mini-*white*, we stabilize both the “ON” and the “OFF” transcriptional states, allowing it to function as well as the transgene with two boundaries. Finally, deleting two imaginal disc enhancers from the *invΔ33* locus led to variegated En expression, suggesting that these enhancers are chromatin regulated and that additional imaginal disc enhancers exist somewhere in this DNA.

### En represses its own expression

Previous studies have shown that overexpression of En can silence the expression of the En gene in wing discs (Tabata et al. 1995; Guillén et al. 1995). Here we show that wildtype levels of Inv and En repress the expression of reporter genes that contain either fragment O or S in the posterior compartment of the wing disc. Furthermore, HA-en expression from the *HAen79* transgene is variegated in the wing pouch of the disc in a wildtype background. Repression of the *HAen45* transgene is even stronger and is relieved by an En point mutation that produces no En protein, showing that the repression is protein-mediated, and not via an interaction of the transgene with the endogenous locus as has been seen for *spineless* (Viets et al. 2019). Removing the imaginal disc enhancers from *HAen79* increases its repression in a wildtype background, suggesting there is a competition between repression and activation. Finally, while we suspect that Inv may also play a role in repressing expression from the *HAen45* and *HAen79* transgene, our results do not directly address this question.

ChIP experiments indicate that the En protein binds to both the O and S fragments, providing evidence that the repression by En may be direct. Removing either fragment ‘O’ or ‘S’ from *HAen79* makes it more susceptible to repression by En. En is also bound to other places in the *inv-en* domain, including the *en* PREs (Fig. 2), and we suggest that En represses itself through many DNA fragments in its endogenous locus. When a gypsy boundary is added to *HAen79*, HA-en expression is no longer variegated in a wildtype background, and Inv, coming from the wildtype *inv-en* domain, has variegated expression. This suggests that En can also repress Inv expression. The Inv and En proteins largely overlap in wing discs, but their expression levels vary, and in the wing hinge, Inv is more highly expressed. The *inv* and *en* promoters are paired in embryos and imaginal discs; these two genes are in the same H3K27me3 domain, and they share many enhancers (Cheng et al. 2014). Therefore, we think of *inv* and *en* as co-regulated genes. However, the levels of expression of Inv and En are not the same, and we suggest that they could directly influence each other’s expression.

### The *invΔ33* domain is more stable than the *HAen79* transgene

To compare the activity of the *HAen79* transgene with the endogenous *inv-en* domain, we deleted 33kb of *inv* DNA, creating the allele we call *invΔ33*. Deletion of fragments O, S, or SS2 from *invΔ33* (*invΔ33ΔO*, *invΔ33ΔS*, *invΔ33ΔSS2* or *HAinvΔ33ΔSS2* heterozygous to *inv-en* deletions) did not lead to any defects in viability. Further, aside from holding their wings out, an indication of loss of *inv* DNA (De et al. 2016), there were no wing defects in these flies. In contrast, deletion of fragments O, S, or SS2 from the *HAen79* transgene led to defects indicative of a loss of *en* function, including defects in wing veins in the posterior compartment and the formation of anterior bristles on the posterior margin of the wing. Flies with a single copy of *HAen79ΔO enΔ110* or *HAen79ΔS enΔ110* did not hatch and had leg defects, indicating that these enhancers are also important for leg development. These data show that the endogenous locus is more resilient than the transgene to deletion of enhancers. Thus, our transgene experiments allowed us to detect enhancer activities that would not have been evident if experiments were only conducted on the endogenous *en* locus.

Deletion of both fragments O and SS2 from *invΔ33* led to variegated expression of En with variable wing defects. This result tells us two things: (1) there are other imaginal disc enhancers located within *invΔ33*, and (2) the variegated pattern suggests that the imaginal disc enhancers are regulated by chromatin. We suggest that there is a competition between the “ON” and “OFF” states. In the presence of all the imaginal disc enhancers, the memory of the “ON” transcription state is maintained in the posterior compartment, perhaps by the trithorax group proteins, and in the “OFF” state, the Polycomb group proteins are the winners of the competition.

### Adding a gypsy boundary to one or both sides of the *HAen79* transgene improves its function

Unlike the endogenous *en* locus, the *HAen79* transgene does not have any boundary elements. We previously showed that the *H3K27me3* domain that forms over this transgene extends into flanking DNA until it is stopped by transcribed genes (De et al. 2019). We also showed that genes flanking the *HAen79* DNA are expressed in a subset of En-expressing cells in embryos. Here we show that the flanking genes mini-*white* and *yellow* are also expressed in wing discs. We hypothesized that adding boundary elements to the ends of *HAen79* would restrict the enhancers to the HA-en gene, improving its function. Surprising to us, adding a gypsy boundary between *HAen79* and the mini-*white* gene had the same effect on its function as adding it to both sides (Fig. 8, Table 1). Both *HAen79GyMW enΔ110* and *HAen79GyB enΔ100* are viable over *inv-gene* deletions with no wing vein defects. They also both make the expression of the HA-en transgene more robust in a wild-type background. Why should insertion of gypsy on only one side work as well as on both sides? We suggest that the enhancers that are chromatin regulated are located upstream of the En transcription unit, and the blocking of these enhancers directs all their activity to the HA-en gene. This is sufficient for complete rescue of the phenotypes we assayed.

An alternative explanation is that the boundary blocks transcription of a transcript initiated at the 5’P element promoter located upstream of *en* in *HAen79* (Fig. 7A). A recent paper showed that transcripts generated by the P-element promoter in the absence of the Homie boundary are highly processive and repress *even skipped* enhancer and promoter activities (Fujioka et al. 2021). In this model, *en* enhancers would stimulate both the expression of both the mini-*white* and the P-element promoter. Read-through transcription from the P-element promoter, through the mini-*white* gene, and perhaps even the *en* regulatory and promoter regions, could interfere with their activity. When gypsy is inserted between *en* and mini-*white*, the P-element promoter is not stimulated in an *en*-like pattern, and read-through transcription does not destabilize *en* expression.

### Similarities between *Ubx* and *en* imaginal disc enhancers

The homeotic gene *Ubx* specifies the formation of the haltere and must be silenced in the wing disc to prevent wing to haltere transformations. Nevertheless, there is an imaginal disc enhancer (IDE) within the *Ubx* locus that, when included in a reporter construct, causes the reporter gene to be expressed in both the haltere and wing discs. Combining this IDE with an embryonic *Ubx* enhancer that sets the boundaries of reporter gene expression in the embryo leads to reporter expression only in the haltere disc (Pirrota et al 1995; Poux et al. 1996; Müller and Bienz 1991; Christen and Bienz 1994). The silencing of the IDE enhancer is due to the Polycomb group genes. As shown here, the En IDEs work in a similar way. Another similarity is that high levels of Ubx protein can silence its own expression (Irvine et al. 1993; Garaulet et al. 2008; Crickmore et al. 2009). In fact, Ubx represses *Ubx* through directly binding to an IDE (Delker et al. 2019). Further, mutations in Ubx binding sites within this IDE caused a loss of repression of a reporter gene but had no detectable effect on *Ubx* expression when made in the endogenous locus (Delker et al. 2019). Thus, *Ubx* autoregulation modulates its own expression level throughout the haltere disc, and En (and Inv) likely does the same in the posterior compartment of wing imaginal discs.

### Concluding Remarks

Inv and En are essential for Drosophila development in both embryos and imaginal discs, and the genome has devoted a lot of DNA to ensure their correct expression. We previously showed that the constitutive PREs are not required for viability or H3K27me3 domain formation at the *inv-en* domain in the laboratory (De et. al. 2016). Here we show that two imaginal disc enhancers are also not required for viability in the laboratory. What is not captured in our papers is that these PRE or enhancer deletion lines are not completely wildtype and are susceptible to mutations elsewhere in the genome. In fact, deleting the PREs from *invΔ33* made it susceptible to a mutation in the Polycomb group gene *ph* (Fig. 8). When we first isolated the *HAen79ΔO* and *HAen79ΔS* transgenes, there was a second site mutation located at another place on the second chromosome that made the wing phenotypes of these flies much more severe in *en* mutant backgrounds. It is possible that *invΔ33ΔO* and *invΔ33ΔS* flies are also susceptible to second site mutations, although we did not test this. Thus, as seen at other loci (reviewed in Kvon et al. 2021), we suggest that the seemingly redundant *inv-en* enhancers and PREs impart a stability that ensures robust development and resiliency important for survival outside of the laboratory.

## Materials and methods

### Small transgenes

Fragments O (2R:7,435,274..7,439,183) and S (2R:7,448,809..7,451,6450 were cloned in a vector that contains about 8kb of upstream *en* regulatory DNA, including the *en* promoter (fragment H, 2R:7415785-7423711, Cheng et al. 2014) (Supplemental Figure 1). All genomic coordinates are in genome release dm5. Fragments S, SS1(2R:7,450,142..7,451,645), and SS2 (2R:7,448,809..7,450,141) were cloned into the vector *pBPGUw* using the procedures described (Pfeiffer et al. 2008).

### CRISPR/Cas9 mutants

The following mutant strains were generated with *CRISPR/Cas9* technology: *enΔ110* (De et al. 2016), *invΔ33* (De et al. 2016), *invΔ33ΔO*, *invΔ33ΔS*, *invΔ33ΔSS2*, *invΔ33ΔOΔSS2*, *HAinvΔ33ΔSS2*, *HAen*, *HAen79GyMW*, and *HAen79GyB* (this paper). Genomic coordinates: *invΔ33*, 2R:7,353,764..7,386,881 (De et al. 2019), ΔO, 2R:7,435,274..7,439,183; ΔSS2, 2R:7,448,809..7,450,141, ΔS in *invΔ33ΔS* is not a simple deletion: it deletes 2 fragments, one is 2683 bp, 2R:7,448,714..7,451,396 that has a 49bp insert sequence of unknown origin at the 3’ end of the deletion, and a smaller 241bp deletion, 2R:7,451,535..7,451,775. Note that 138bp between these two fragments is present.

For CRISPR target sequence design and cloning into the pU6-BbsI-gRNA(chiRNA) vector we followed the protocols in https://flycrispr.org. Repair plasmids were used to make all new mutant fly lines except *invΔ33ΔS*. The repair plasmids were either synthesized by Genscript Inc or by assembling PCR fragments with NEBuilder (New England Biolabs). Typical repair plasmids had homology arms of 500 bp to 1000bp, depending on the experiment. The cloning vector for repair plasmids was pUC57.

The gRNA plasmids were mixed equally to get a total concentration of 1 µg/µl. When a repair plasmid was used, the repair plasmids were mixed with gRNA plasmids at 0.5 µg/µl. The plasmid mixture was injected (Rainbow Flies, Inc or BestGene, Inc) into the relevant host fly strain expressing Vasa-Cas9 (Sebo et al. 2014). Genotypes injected were *M{vas-Cas9.S}ZH-2A*; with an appropriate 2^nd^ chromosome: wildtype, *invΔ33*, *invΔ33ΔSS2*, *HAen79*, or *HAen79GyMW*. Adult flies from the injected embryos (G0) were singly crossed to a stock with a second chromosome balancer, *yw; Sco/CyO*. After about a week at 25°, when larval progeny were present, DNA was prepared from single fertile G0 flies. PCR was done to detect the desired modification in the single G0 flies. From G0s that gave a strong PCR signal, 20 of its progeny (G1) were singly crossed to *yw; Sco/CyO*. PCR was done on each G1 fly. The G1 flies that gave the right PCR product were used to establish a fly stock. After the fly stock was established, PCR and DNA sequencing were done to verify the desired change. A more detailed *CRISPR/Cas9* protocol for the generation of *HAen79GyMW* is described below. For other experiments, detailed protocols are available upon request.

The fly strain *y^1^ M{vas-cas9.S}ZH-2A w^1118^; HAen79* was generated using standard genetic crosses. A gRNA target site, ggccggccgcgatcgcgccc, was identified between mini*-white* gene and the 5’ end of *en* DNA present in *HAen79*. A repair plasmid was made using NE Builder (New England Biolabs) that includes a 430bp gypsy insulator sequence (Geyer and Corces 1992) flanked by two 1kb homology arms. G0 and G1 flies were screened and a fly stock was established. PCR and DNA sequencing were done to verify the gypsy insertion and flanking sequences, resulting in the *HAen79GyMW* strain.

### Tagging en with HA

A 27 bp DNA sequence, TACCCCTACGACGTCCCCGATTACGCC, that encodes the 9 amino acids HA tag was inserted into *en* coding sequence immediately downstream of its translation start codon ATG onto either a wildtype second chromosome (*HAen*) or *invΔ33ΔSS2* (*HAinvΔ33ΔSS2*).

### Immunostaining and RNA-FISH

Our antibody staining procedure for imaginal discs has been described previously (Cheng et al. 2014). The primary antibodies used were: guinea pig anti-Inv (1:5000, Cheng et al, 1014), rabbit anti-EN (1:500, Santa Cruz Biotechnology, Inc.), anti-HA.11 (1:1000, clone 16B12, Biolegend), rabbit anti-Gal4 activation domain (1:1000, Millipore Sigma). Alexa Fluor secondary antibodies were used (1:1000, Invitrogen) and discs were mounted in Vectashield with DAPI (Vector Labs). Fluorescent RNA in situ hybridization was done with DIG-labeled probes for *white* and *yellow* with TSA Plus Fluorescence Kits (Akoya Biosciences) and used for in situ hybridization to imaginal discs as described (Tian et al. 2019).

### Large transgenes

Generation of the *HAen45*, *HAen79*, and *HAinv84* transgenes and transgenic lines were previous described (Cheng et al, 2014). The transgenes *HAen79ΔS*, *HAen79ΔO* and *HAen79ΔSS2* were generated using recombineering to delete DNA using the *HAen79* plasmid as the starting construct (for procedures see Cheng et al. 2014). Detailed protocols are available upon request. The coordinates of the deletions were the same as in *invΔ33*, except that ΔS is a simple deletion in *HAen79ΔS* (2R: 2R:7,448,809..7,451,645). These transgenes were inserted at the attP40 insertion site (on chr2L) and recombined with *enΔ110*, *en^E^*, or *en^X31^* (on chr2R) to generate the chromosomes used to test the transgene’s function in the absence of endogenous *inv* and *en*. PCR was used to detect the presence of the transgene and the *en* mutations.

## Acknowledgements

We thank Drs. Pedro Rocha, James Kennison, James Jaynes, and Lesley Brown for helpful comments on this manuscript.

---

**Fig. S1.**
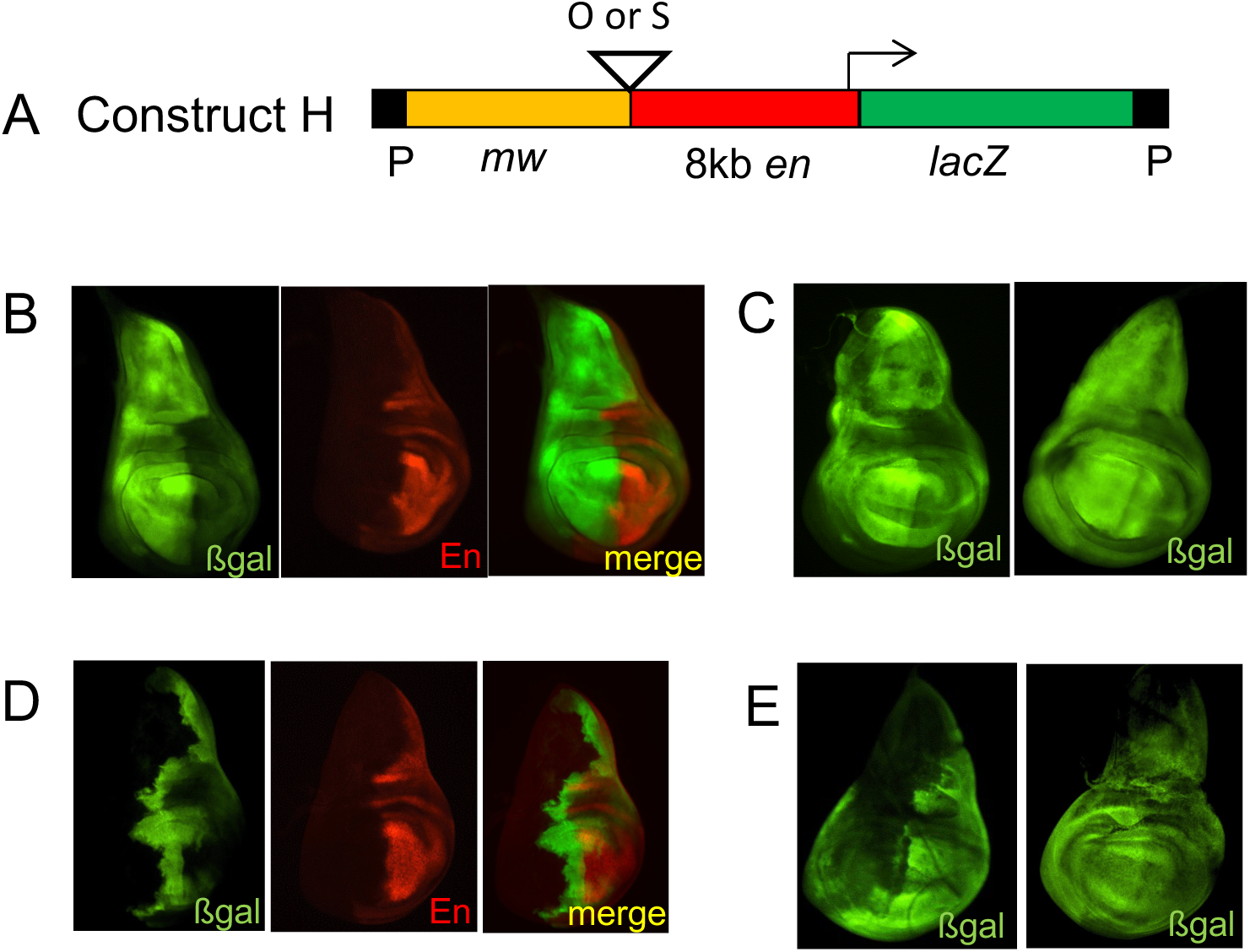
Activity of enhancers ‘O’ and ‘S’ in reporter constructs. (A) Diagram of *en-lacZ* reporter construct used. The 8kb *en* fragment drives *lacZ* expression in embryos in stripes but there is no expression of *lacZ* in the imaginal discs (Fragment H in Cheng et al 2014). (B, C) Expression of ßgal and En (B) and ßgal (C) from *S-enlacZ* inserted at 3 different insertion sites. (D,E) *O-enlacZ* inserted at 3 different insertion sites.

**Fig. S2.**
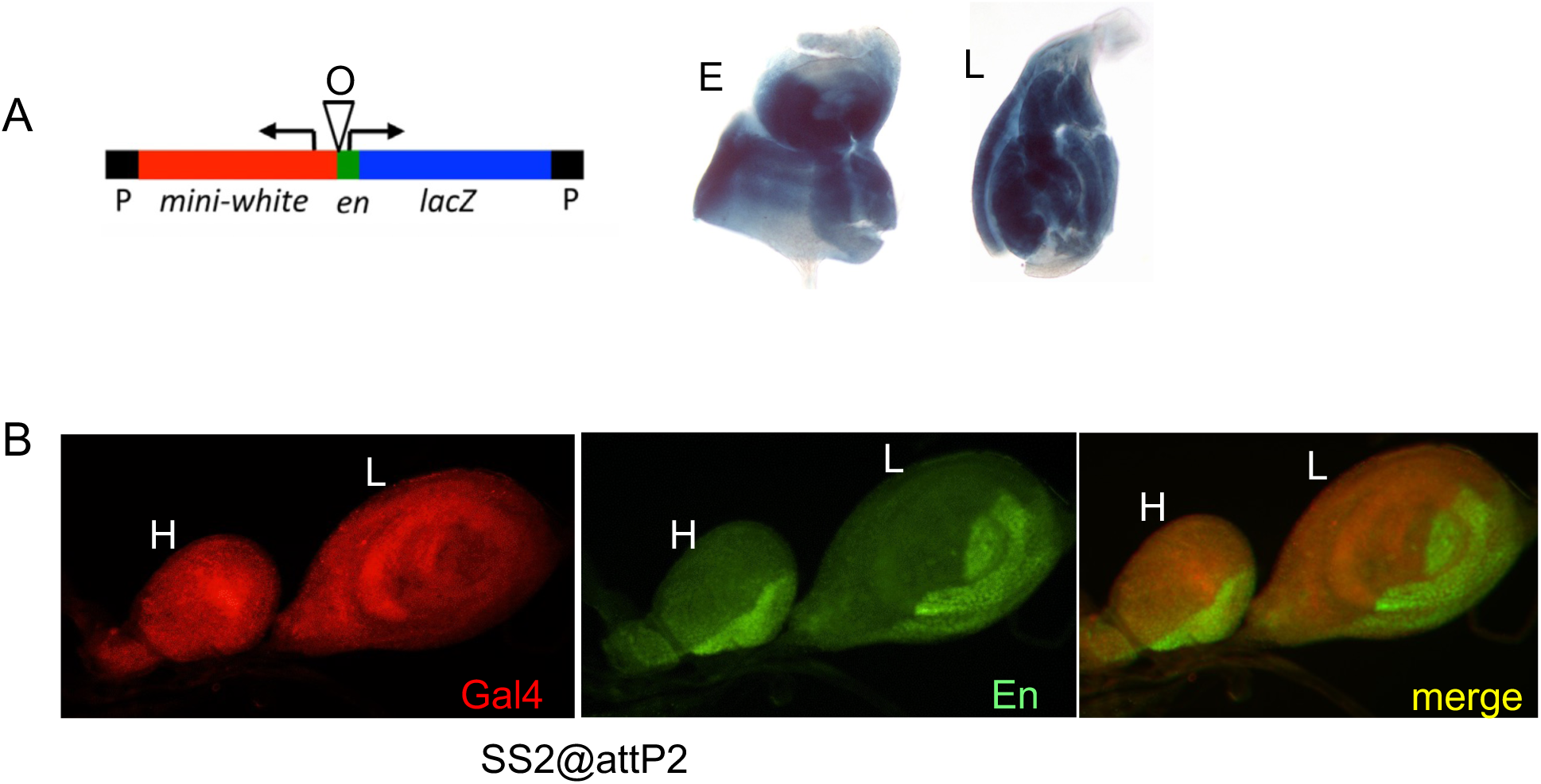
Fragments O and S drive expression in other discs. (A) DNA construct and ßgal staining of imaginal discs from third instar larvae showing that the O enhancer drives expression of lacZ in the eye (E) and leg (L) discs (construct from Cheng et al. 2014). (B) Gal4 expression from SS2@attP2 in the haltere (H) and third leg (L) disc. There is less expression of Gal4 in the posterior part of the leg disc. In this halter disc, Gal4 appear to be expressed in the entire disc but in other haltere discs it was reduced in the posterior compartment.

